# Potassium Carbonate (K_2_CO_3_) – A cheap, non-toxic and high-density floating solution for microplastic isolation from beach sediments

**DOI:** 10.1101/2020.11.17.386417

**Authors:** Jan Gohla, Sandra Bračun, Gerwin Gretschel, Stephan Koblmüller, Wagner Maximilian, Christian Pacher

## Abstract

Beaches are good indicators for microplastic distribution and local microplastic pollution. Multiple methods have been developed for extracting microplastics from sediment, mainly through density separation. However, the chemicals applied are often expensive and harmful for the user or to the environment. We briefly review the problems associated with the use of these chemicals and present a new floatation solution, potassium carbonate (K_2_CO_3_) that has many advantages over available media. It is non-toxic and cheap, and with a density of 1.54 g/cm^3^ the K_2_CO_3_ solution yielded a mean recovery rate of around 92% for PVC, one of the densest polymers, that cannot be easily extracted with alternative floatation agents. We propose that the use of K_2_CO_3_ is particularly promising for long term and large-scale monitoring studies, not least because it allows an increasing involvement of citizen scientists, hopefully leading to an increased public awareness of the plastic problem in the seas.

---

The impact and ubiquity of plastics in the environment has become pervasive enough that it has been proposed as a stratigraphic indicator for the Anthropocene (Zalasiewicz et al., 2016). Five huge plastic vortices, containing all kinds of garbage, have been reported from our oceans (Eriksen et al., 2013; Law et al., 2010; Lebreton et al., 2018; Van Sebille et al., 2012). Yet, more than 92% of the plastic waste in the sea consists of particles smaller than ca. 5 mm, so-called microplastic (Eriksen et al., 2014). In addition to often incorporated environmentally damaging additives (e.g. softeners like phthalates which are used to increase the flexibility of polymers), other poisonous substances are prone to adhere to microplastics and, if ingested by organisms, these can be accumulated in the food web (e.g. Anastasopoulou et al., 2013; Avio et al., 2015; Farrell and Nelson, 2013; Fossi et al., 2017; Rochman, 2015; Romeo et al., 2015; Setälä et al., 2014).

The presence of microplastic has been confirmed in the water column and marine sediments throughout the world (e.g. Claessens et al., 2011; Law et al., 2010; Moore et al., 2001; Thompson et al., 2004). Mean densities of most common plastic types range between 0.9 and 1.5 g/cm^3^. Particles with a density lower than seawater (<1.02 g/cm^3^) will float on the sea surface or drift in the water column and those with a higher density will sink to the seafloor and accumulate in the sediment. Besides visual and manual sorting of larger particles, the most common method for extracting microplastic from beach sediments is density separation (e.g. Hidalgo-Ruz et al., 2012; Van Cauwenberghe et al. 2015, Mahat, 2017); i.e., all polymers with a lower (or theoretically equal) density than the floatation solution can be separated from the inorganic substrate by suspension. However, multiple difficulties arise, especially considering efficiency, toxicity or economic feasibility of available floating reagents (Fig. 1). In order to properly quantify the amount of microplastic pollutants in the environment, floatation agents are needed that cover the entire density range and do not decompose or change the molecular or physical structure of the polymers. Additional problems appear through the so called “clumping-effect”, which occurs when dissolved organic matter (e.g. proteins or polysaccharides) aggregates around sediment particles. Owing to this, it becomes difficult or impossible to extract “clumped” (i.e. high density) microplastic particles from the sediments by density separation.

**Fig. 1.**
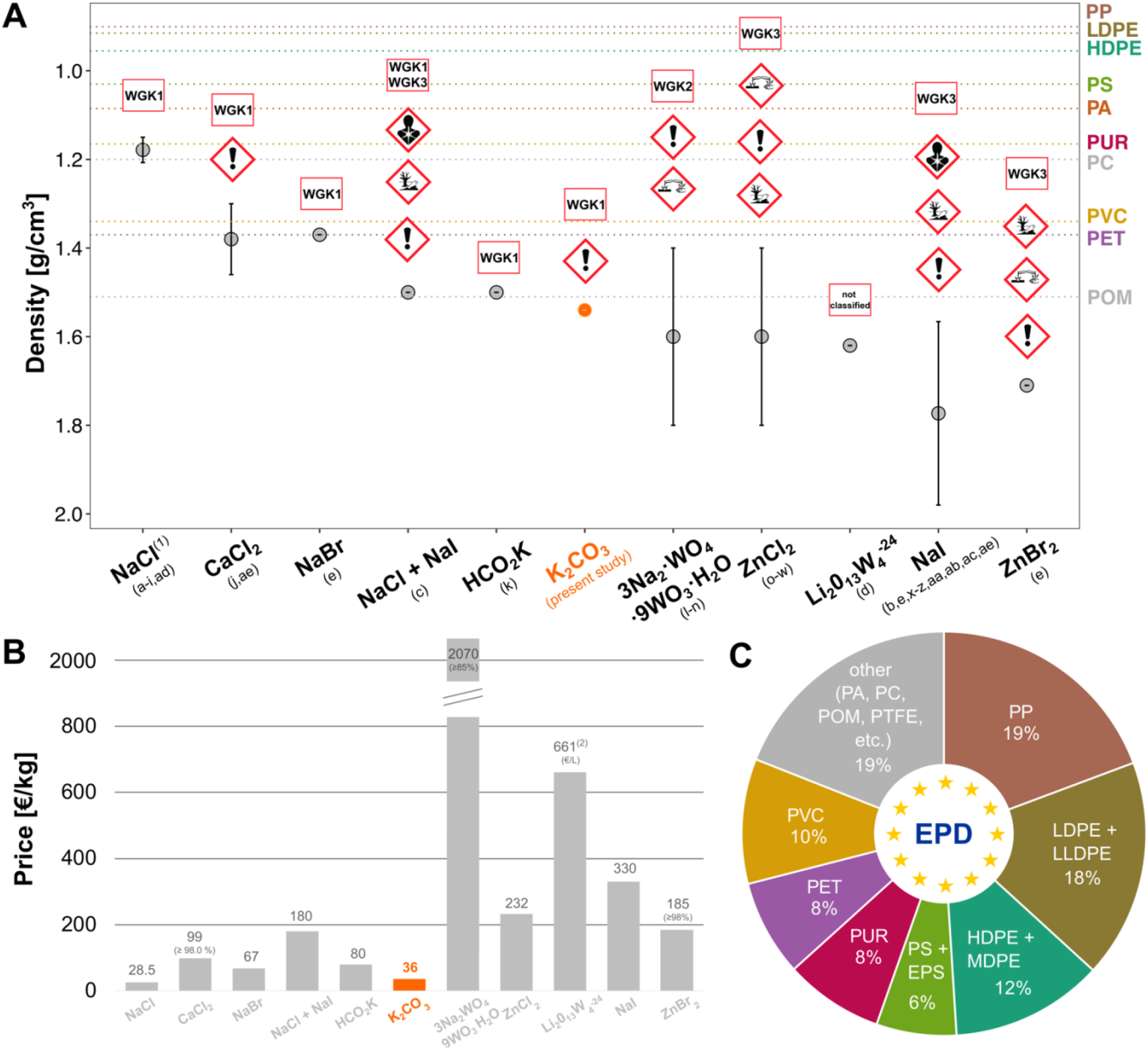
Overview of efficiency and toxicity of different microplastic extraction flotation solutions. **(A)** Dots show mean measured density [g/cm^3^] of commonly used microplastic flotation solutions. If applicable, ranges of densities used in different studies (a-z and aa-ae; see below) are given. Dashed coloured horizontal lines indicate mean densities of different common polymers (excluding EPS and PTFE), obtained from different sources (Hamm et al., 2018; Han et al., 2019; Hidalgo-Ruz et al., 2012; Nuelle et al., 2014; Qiu et al., 2016; Quinn et al., 2017; US EPA, 1992). Note that the y-axis is switched in order to visualize the “floating” capacity (i.e. lower density floats on higher density). GHS-Symbols (Globally Harmonized System of Classification, Labelling and Packaging of Chemicals) retrieved from Sigma Aldrich, product data sheets. WGK (“Wassergefährdungsklasse” = Water Hazard Classification) retrieved from Carl Roth, product data sheets and Umweltbundesamt IV 2.6. (B) Relative costs [€/kg] of different floatation methods (99% purity) obtained from https://www.sigmaaldrich.com/catalog (accessed: 10.13.20). **(C)** European plastic demand (EPD) in [%] (PlasticsEurope, 2019). Abbreviations: Sodium chloride (NaCl), Calcium chloride (CaCl_2_), Sodium bromide (NaBr), Potassium formate (HCO_2_K), Potassium carbonate (K_2_CO_3_), Sodium polytungstate (3Na_2_WO_4_·9WO_3_·H_2_O), Zinc chloride (ZnCl_2_), Lithium metatungstate (Li_2_O_13_W_4_^-24^), Sodium iodide (NaI), Zinc bromide (ZnBr_2_); Polypropylene (PP), Low-Density Polyethylene (LDPE), Linear Low-Density Polyethylene (LLDPE) High-Density Polyethylene (HDPE), Medium-Density Polyethylene (MDPE), Polystyrene (PS), Expanded Polystyrene (EPS), Polyamide (PA), Polycarbonate (PC), Polyurethane (PUR), Polyethylene terephthalate (PET), Polyvinyl chloride (PVC), Polyoxymethylene (POM), Polytetrafluoroethylene (PTFE); low hazardous to waters (WGK1), distinctly hazardous to waters (WGK2), severely hazardous to waters (WGK3); Sources: ^a^Carson et al., 2011; ^b^Dekiff et al., 2014; ^c^Han et al., 2019; ^d^Masura et al., 2015; ^e^Quinn et al., 2017; ^f^Reddy et al., 2006; ^g^Thompson et al., 2004; ^h^Vianello et al., 2013; ^I^Yu et al., 2016; ^j^Stolte et al., 2015; ^k^Zhang et al., 2017; ^l^Ballent et al., 2016; ^m^Corcoran et al., 2009; ^n^Enders et al., 2020; °Bergmann et al., 2017; ^p^Imhof et al., 2012; ^q^Imhof et al., 2013; ^r^Imhof et al., 2016; ^s^Imhof et al., 2018; ^t^Liebezeit and Dubaish, 2012; ^u^Löder et al., 2017; ^v^Mintenig et al., 2017; ^w^Rowe et al., 2019; ^x^Van Cauwenberghe et al., 2013a; ^y^Van Cauwenberghe et al., 2013b; ^z^Claessens et al., 2013; ^aa^Nuelle et al., 2014; ^ab^Kedzierski et al., 2017; ^ac^Fischer and Scholz-Böttcher, 2017; ^ad^Claessens et al., 2011; ^ae^Crichton et al., 2017; ^1^Yu et al. (2016) stated a density of 1,27g/cm^3^, which is not possible for NaCl-Solution, hence a typing error was assumed and the highest possible density at 0°C was put instead (retrieved from https://handymath.com/cgi-bin/nacltble.cgi?submit=Entry; 2020-04-16). ^2^Price retrieved from: https://www.lmtliquid.com/price-and-order-information.html; 2020-09-21.

Other commonly used floatation agents are often expensive or harmful to the environment and the user (Claessens et al., 2013; Han et al., 2019; Kedzierski et al., 2017; Miller et al., 2017; Nuelle et al., 2014). A commonly used non-toxic and cheap floatation agent is a saturated sodium chloride (NaCl) solution (e.g. Thompson et al., 2004). However, plastics with a density greater than 1.2 g/cm^3^, like polyvinyl chloride (PVC) or polyethylene terephthalate (PET), cannot be extracted using this agent. Nonetheless, these polymers make up almost one fifth of the European plastic demand (Fig. 1C; PlasticsEurope, 2019), are present in large quantities in textile fibres and drinking bottles (Coppock et al., 2017) and are the first to be incorporated into marine sediments (Van Cauwenberghe et al., 2015). Hence, the use of NaCl highly underestimates and neglects considerable amounts of microplastics in marine sediments. In contrast, high density floating agents, such as zinc chloride (ZnCl2) (e.g. Imhof et al., 2013; Liebezeit and Dubaish, 2012), sodium iodide (NaI) (e.g. Dekiff et al., 2014; Van Cauwenberghe et al., 2013a; Van Cauwenberghe et al., 2013b), or sodium polytungstate (3Na2WO4·9WO3·H2O) (e.g. Ballent et al., 2016; Corcoran et al., 2009; Enders et al., 2020), are capable of extracting PVC or PET. Due to their tremendous cost, toxicity and environmentally harmful nature, the handling of these substances requires lots of experience and cautiousness. For instance, ZnCl2 is explicitly mentioned to be very toxic to aquatic life, with long lasting effects (GHS hazard statement: H410), NaI is declared as severely hazardous to water systems (since 2018 WGK3 according to the environmental agency Germany (=Umweltbundesamt) IV 2.6) and very toxic to aquatic life (H400), sodium polytungstate is distinctly hazardous to waters (WGK2) and harmful to aquatic life with long lasting effects (H412) and zinc bromide is toxic to aquatic life with long lasting effects (H411) (Fig. 1; Sigma Aldrich and Carl Roth, product data sheets). Altogether, this makes the application of these solutions impractical and inconvenient for large scale studies, surveys without access to chemistry laboratories or research that involves non-experts, such as citizen science projects (Lots et al., 2017; Rambonnet et al. 2019; Carbery et al., 2020).

Here, we present a new, cheap, non-toxic and high-density floatation solution for isolating microplastics from beach sediments, potassium carbonate (K_2_CO_3_). We elaborate that potassium carbonate obtains the necessary properties to extract – with a high rate of recovery – polymers with a density of up to 1.5 g/cm^3^. This covers the most abundant plastics in Mediterranean beach sediments such as PE, PP, PS, PET, and PVC (Digka et al., 2018) and lays the foundation for new applications in the field of microplastic research and large-scale monitoring as well as conservation projects. There are several advantages of potassium carbonate (K_2_CO_3_) over currently used common floatation solutions, especially regarding its environmental sustainability, toxicity, physiochemical properties and costs. According to the Globally Harmonized System of Classification and Labelling of Chemicals (GHS), potassium carbonate is less dangerous and environmentally harmful than most commonly used floatation media (Fig. 1). For instance, in comparison to zinc chloride, a heavy metal solution, which after usage has to be collected and disposed of at great financial and material cost, K_2_CO_3_ can be disposed without harming the environment (WGK1; Fig. 1). Above all, during handling, potassium carbonate is equivalent to the hazardous material classification of commercial dish washing soap (Fig. 1).

Potassium carbonate reaches densities high enough (max. approx. 1.8 g/cm^3^; Fig. 1B) to allow the suspension of nearly all common sorts of plastics in beach sediments (in this study a density of 1.54 g/cm^3^ was used; Fig. 1). Furthermore, the alkaline nature (pH 11.5) of K_2_CO_3_ amplifies the dissolving process of biological organic components through hydrolysis. Consequently, compared to other floatation solutions, K_2_CO_3_ tremendously reduces the mechanical effort (i.e. stirring) needed to overcome the above mentioned “clumping effect”, increases the overall efficiency and allows the analysis of more samples per time (own observation). Previous (Bürkle GmbH, 2020) as well as own visual investigation of plastic particles after K_2_CO_3_ treatment have shown that plastics – with the exception of polycarbonate (PC) (Bürkle GmbH, 2020) – are chemically resistant towards potassium carbonate. However, since PC only makes up around 2% of yearly produced plastics in Europe, the relative role of PC in shore sediments can be neglected (Fig. 1C; PlasticsEurope, 2019). Finally, in comparison to other floatation solutions (Fig. 1B), potassium carbonate is cheap (36.20 €/kg; but can also be found cheaper for 5.65 €/kg online) and easily accessible, a prerequisite to conduct high throughput studies including lots of samples.

Handling, preparing and storing a ready-to-use potassium carbonate is simple and fast (Fig. 2). K_2_CO_3_ powder should be stored in a dry, closed place at room temperature (e.g. sealable bucket or glass bottle). To obtain a solution of 1.54 g/cm^3^ (efficient enough for most polymers), 770 g of K_2_CO_3_ (in small portions) are dissolved under constant stirring in 500 - 600 ml dH2O and filled up to 1 litre. After 20 minutes of stirring the solution gets clear and heats up due to the exothermic nature of the solvation process. Before usage, the solution has to be cooled down to room temperature and the final density should be measured, by weighing a defined volume of the solution (i.e. 100 ml of the solution need to weigh 154 g, to validate the desired density). The ready-to-use solution can be stored in tinted glass bottles at room temperature for months without any observed negative effects on experiments. The fresh solution sometimes tends to crystalize as dihydrate salt that can easily be resolved by vigorous shaking or using a magnetic stirrer and already used potassium carbonate can be easily recycled, collected and filtered using a vacuum pump and filter-paper (e.g. 4 μm pore size) to avoid left-over particles. The recycled potassium carbonate solution changes its colour from initially colourless to yellow but remains transparent. After 24 hours the recycled solution starts to form streaks at ground level that can be again resolved by shaking or magnetic stirring. Similar to preceding studies that recycled their respective floatation media (Enders et al., 2020; Han et al., 2019; Kedzierski et al., 2017; Nuelle et al., 2014), no impact on the efficiency or any contamination was observed.

**Fig. 2.**
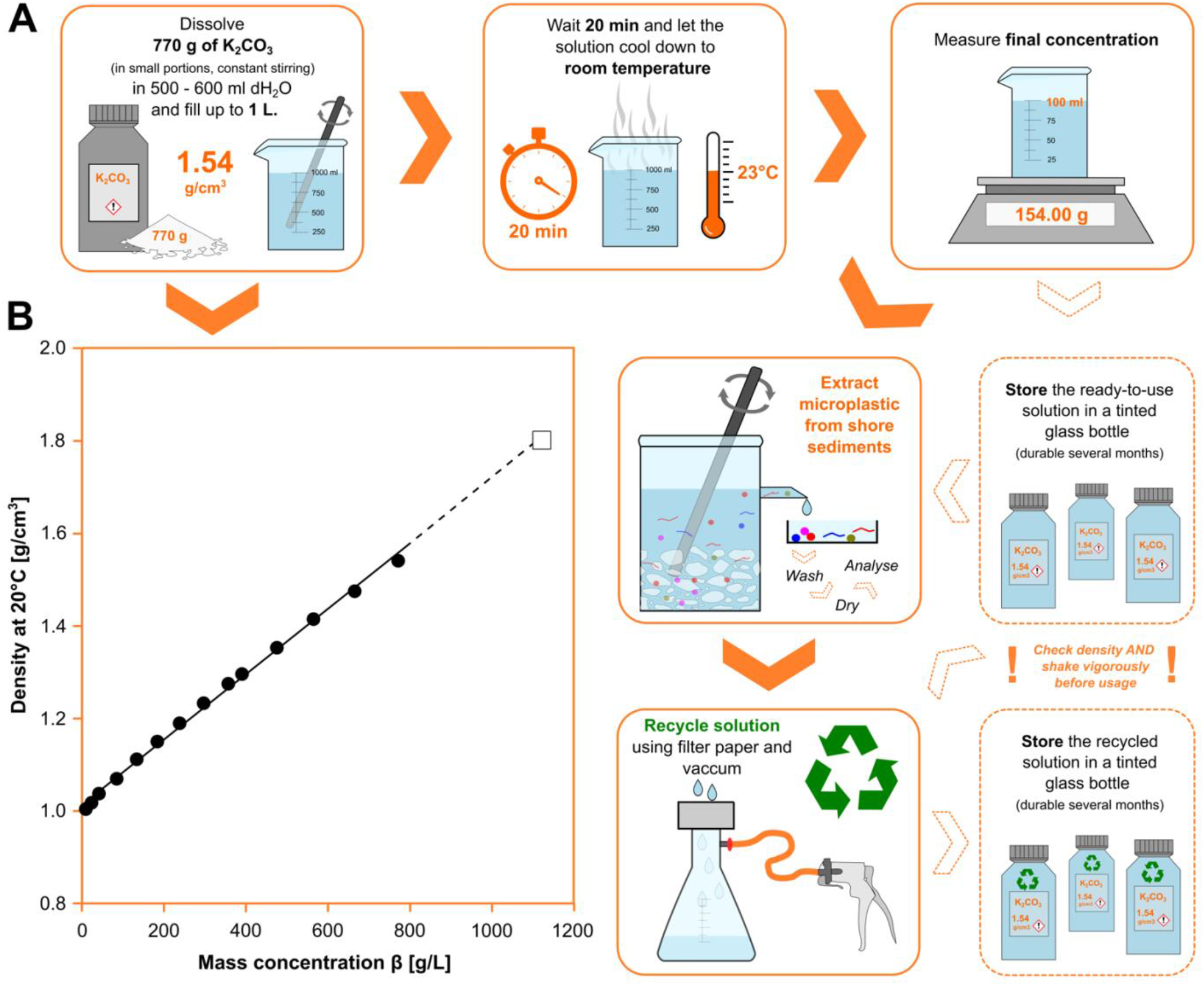
Preparation of a potassium carbonate (K_2_CO_3_) solution for microplastic extraction from beach sediments. **(A)** General workflow for preparing and storage of the ready-to-use K_2_CO_3_ solution with a density of 1.54 g/cm^3^. **(B)** Density of aqueous potassium carbonate solutions as a function of concentration at 20°C (293.15 K). The dotted line is interpolated and ends at 1120 g/L (i.e. the maximum solubility of K_2_CO_3_). Data retrieved from Perry et al. (1997).

To test the efficiency of potassium carbonate, an experimental series using different grain sizes (63μm - 2000μm) and sediment types (cleaned natural beach sediment and commercially available aquarium quartz sand) was conducted using artificially produced polyvinyl chloride (PVC) particles (0.05 g and 0.5 g; Table 1). PVC makes up to ten percent of the European plastic demand and is a common component in beach sediments (Fig. 1C). Its relatively high density (1.1 - 1.58 g/cm^3^) poses problems for density extraction, especially with low density media, such as NaCl. However, K_2_CO_3_ yielded a mean recovery rate of 91.76% (s.d. 8.07%) for PVC, which is comparable to other previously available high-density solutions (reviewed by Miller et al., 2017). For example, the recovery rate for PVC in a comparable particle size class for NaBr yielded around 85% recovery (Quinn et al., 2017). Studies using NaI achieved recovery rates around 97% for PVC, but only using repeated extractions (Crichton et al., 2017) or implementing a two-step procedure including air-induced overflow (Nuelle et al., 2014). Other studies noted a recovery rate of around 90% for PVC in the size category 800-1000 μm (Quinn et al., 2017). ZnBr2 and ZnCl2 solution and PVC particles obtained high recoveries of maximum 95% (Quinn et al., 2017) and 99.7% (particle size 1-5 mm) (Imhof et al., 2012) respectively, whereas the latter ones where extracted using the Munich Plastic Sediment Separator (MPSS), which minimizes particle loss during handling. Without the use of the MPSS, recovery rates were 99,1% for particles sized 1-5 mm and 39,8% for particles smaller than 1 mm. Non-toxic NaCl revealed recovery rates between 68.8% and 97.5% in repeated extractions (Claessens et al., 2011). However, the recoveries depend on particle size and polymer type, whereas larger particles (i.e. 800-1000 μm - comparable to our study) resulted in values below 85% and PVC as well as PET only showed a recovery of 65-70% (Quinn et al., 2017). Nonetheless, due to methodological differences, comparability between different extraction media remains difficult.

**Table 1.** Ranges of the recovery rate and mean recovery rate obtained from five independent replicates with Potassium carbonate (K_2_CO_3_) as floating agent (with a density of 1.54 g/cm^3^) for PVC.

| Artificially added amount of PVC particles | Range of recovery rates in % obtained from 5 replicates |  | Overall mean recovery rate |
| --- | --- | --- | --- |
|  | Natural beach sediment 500 - 2000 µm | Aquarium quartz<sup>1</sup> 63 - 2000 µm |  |
| 0.5 g | 91.31 - 96.47% | 89.76 - 96.76% | **91.76% (s.d. 8.07%)<sup>3</sup>** |
| 0.05 g | 66.73 - 91.09% | 86.92 - 105.40%<sup>2</sup> |  |

As new sampling, extraction and detection methods and techniques are being developed worldwide (e.g. Claessens et al., 2013; Harrison et al., 2012; Imhof et al., 2012; Nuelle et al., 2014), it is clear that, in order to completely understand the distribution of microplastics in the marine environment, a harmonization and standardization of techniques and protocols is urgently needed (Besley et al., 2017). An increased sampling and monitoring effort across large spatial and temporal scales can be achieved by including non-specialists. Several large-scale studies successfully engaged citizen scientists for monitoring microplastic contamination in beach sediments, but these were confined to either direct visual collecting and sorting of particles and/or the use NaCl solutions as for density separation (e.g. Lots et al. 2017; Rambonnet et al., 2019; Carbery et al., 2020). Such studies are infeasible for other floatation media than NaCl, particularly due to tremendous costs that result from large sample sizes. Hence, the use of potassium carbonate could overcome this obstacle and above all increase the efficiency beyond the density threshold achieved by NaCl (Fig. 1). Additionally, the hazard-free workflow (Fig. 2) and trouble-free disposal could allow volunteers to be engaged more in the scientific process. Previous citizen science projects were confined to simple contributions (i.e. sampling/data acquisition only). However, more collaborative (i.e. sample/data analyses) or even co-creative (i.e. set up hypotheses, methods, etc. supervised by trained scientists) frameworks could raise awareness for scientific practice, beyond the underlying environmental problem (Bonney et al., 2009). We, therefore, propose that K_2_CO_3_ bears enormous potential for future microplastic research and long-term conservation and monitoring studies.

## Acknowledgements

For field support and the implementation of the present study we thank the staff of the Morska Škola Pula (Croatia). Furthermore, we thank the Haus des Meeres (Vienna) for supporting this study with the Hans and Lotte Hass prize.

## Notes

### Competing Interest Statement

The authors have declared no competing interest.

